## Supplemental Figures for "Altered auditory feature discrimination in a rat model of Fragile X Syndrome"

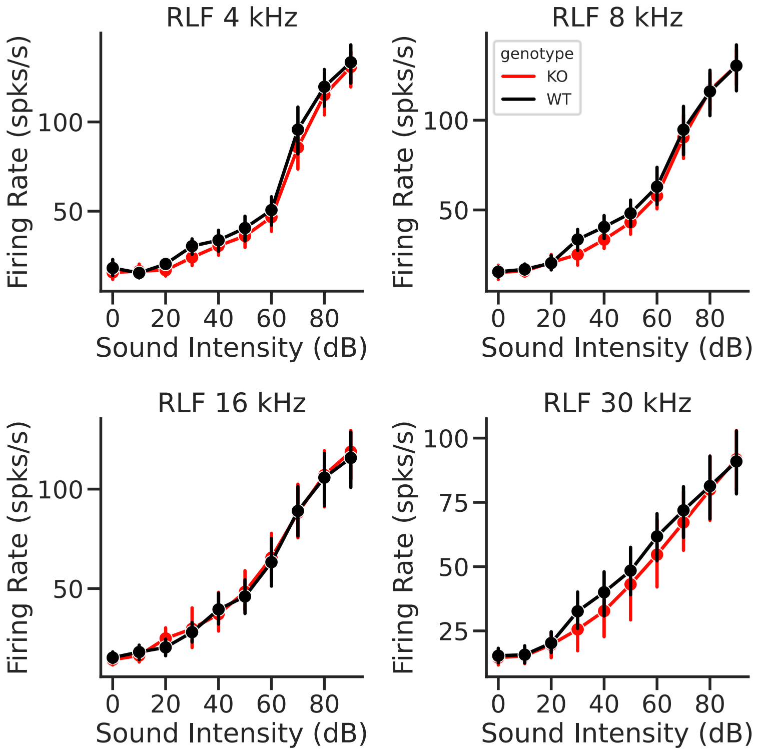


**Supplemental Figure 1. Tone specific rate-levels functions from the inferior colliculus of wildtype and *Fmr1* KO rats.**

Multi-unit spiking activity recorded from the inferior colliculus of wildtype (black) and *Fmr1* KO (red) littermates in response to individual frequencies (4,8,16, and 30 kHz) across intensities (0-90 dB SPL, 10 dB steps). No significant genotype difference was observed at any frequency (GLLM: 4kHz: $p = 0.402$, 8kHz: $p = 0.577$, 16kHz: $p = 0.248$, 32kHz: $p = 0.843$).

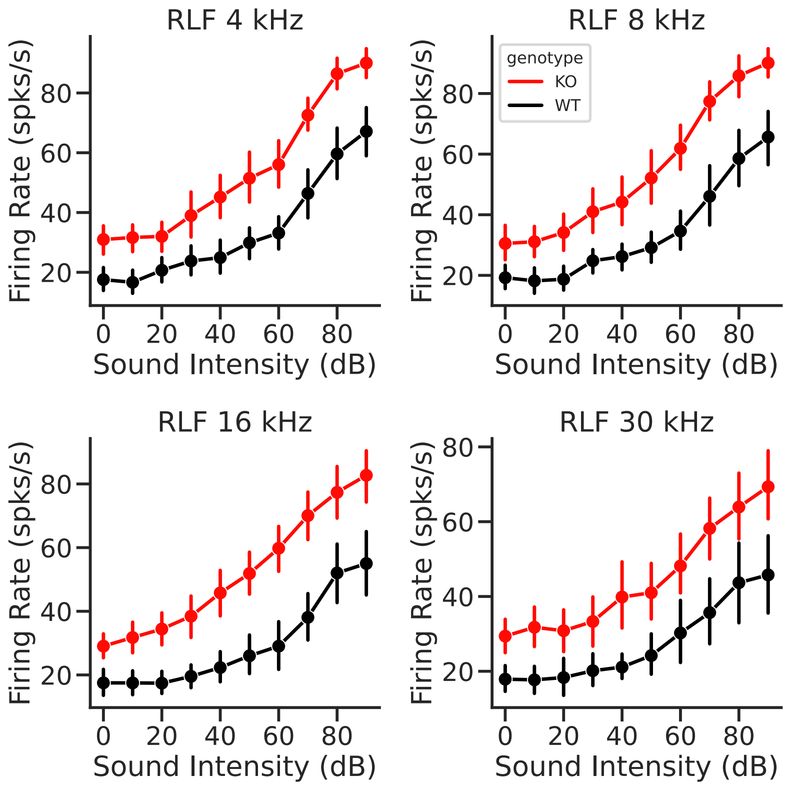


**Supplemental Figure 2. Tone specific rate-level functions from the auditory cortex of wildtype and *Fmr1* KO rats**

Multi-unit spiking activity recorded from the auditory cortex of wildtype (black) and *Fmr1* KO (red) littermates in response to individual frequencies (4,8,16, and 30 kHz) across intensities (0-90 dB SPL, 10 dB steps). Significant genotype differences were observed at each individual frequency (GLMM: 4kHz: *p = 0.012, 8kHz: *p = 0.014, 16kHz: **p = 0.001, 32kHz: *p = 0.024).

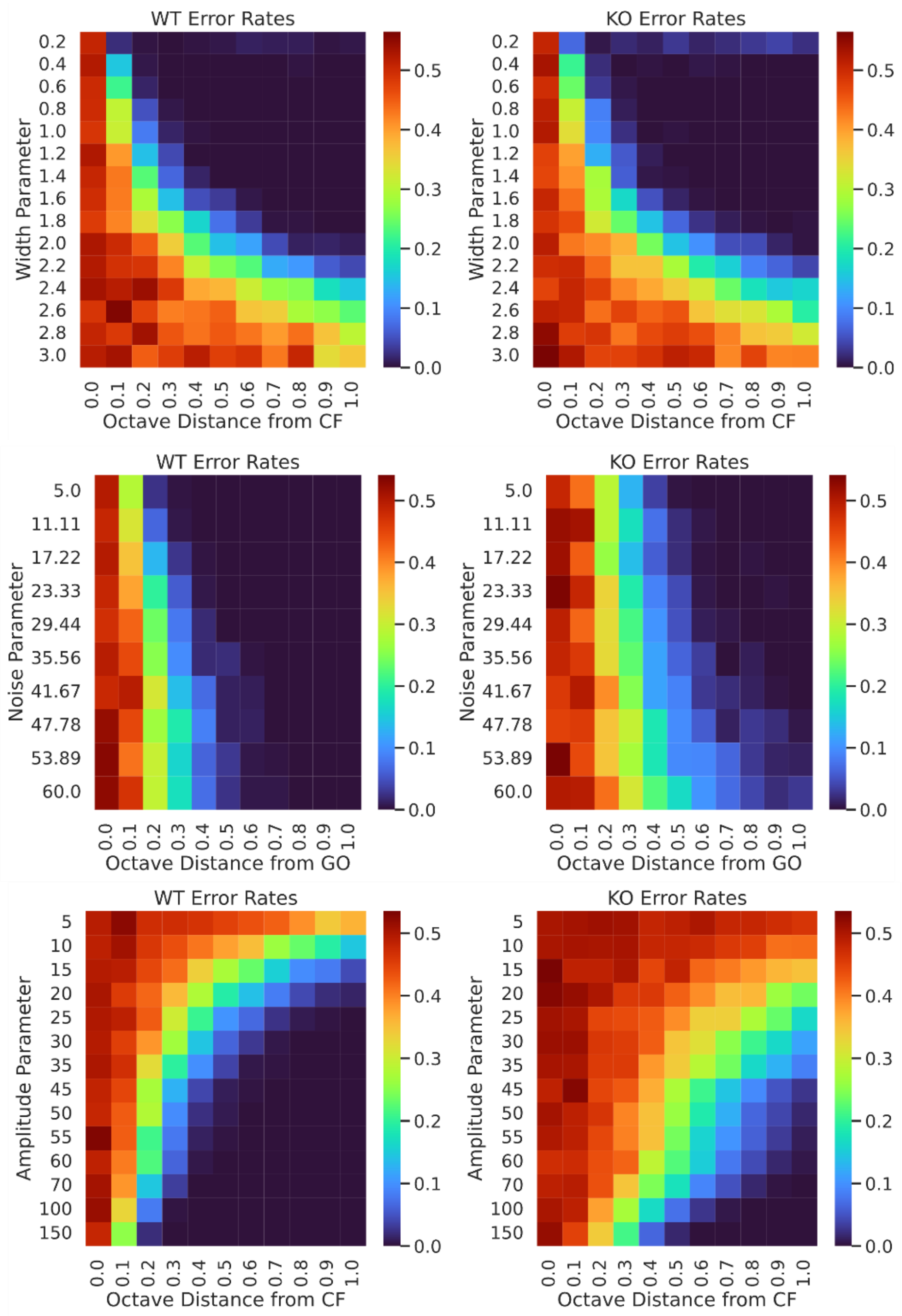


**Supplemental Figure 3. Effect of varying ACx model parameters on decoding accuracy**
**(A)** Heatmaps for error rate as a function of systematically varying the tuning width (for the WT (left) and *Fmr1* KO (right) models. Both KO and WT models appear to be equally sensitive to increased tuning bandwidth. **(B)** Heatmaps for error rate as a function of systematically varying the noise parameter (reflecting spontaneous firing rates) for the WT (left) and *Fmr1* KO (right) models. The KO model appears to be more sensitive to increases in compared to WT. **(C)** Heatmaps for error rate as a function of systematically varying the amplitude parameter (reflecting peak sound-evoked firing rates) for the WT (left) and *Fmr1* KO (right) models. Both model appears are sensitive to reduction in amplitude compared, but KO model has consistently poorer accuracy at all amplitude levels.
